# Reproducible chemostat cultures to eliminate eukaryotic viruses from fecal transplant material

**DOI:** 10.1101/2023.03.15.529189

**Authors:** Signe Adamberg, Torben Sølbeck Rasmussen, Sabina Brigitte Larsen, Xiaotian Mao, Dennis Sandris Nielsen, Kaarel Adamberg

## Abstract

The effect of fecal microbiota transplantation (FMT) on various gut-related diseases is intensively investigated in clinical trials. In addition to bacteria, the gut microbiome also contains eukaryotic, archaeal, and bacterial viruses (bacteriophages, in short phages), which collectively is referred to as the gut virome. Application of FMT in clinical settings is associated with a potential risk for the recipient of transferring infectious eukaryotic viruses or bacteria, despite strict screening procedures for the donor material. A safer and more targeted method to modulate the gut microbiota is therefore needed to extend the application width of FMT. Emerging evidence suggests that gut phages play a key role in maintaining a balanced gut microbiome as well as in FMT efficacy. Thus, a phageome from a cultured fecal donor microbiome may be a more efficient alternative to modulate the gut bacteriome than FMT. Here, we analyzed the dynamic changes of the viromes of mice cecal and human fecal matter inoculated chemostat cultures. Sequencing results showed that the relative abundance of eukaryotic viruses remarkably decreased during continuous cultivation, likely due to the lack of eukaryotic hosts. The corresponding phageome profiles showed dilution rate dependency, a reproducibility between biological replicates, and maintained high diversity of phages although being different from the inoculum phageome. This proof-of-concept study may constitute the first step of developing therapeutic tools to target a broad spectrum of gut-related diseases and thereby replacing FMT with a safer phage-mediated therapy.

## INTRODUCTION

Transplantation of fecal microbiota (FMT) has been successfully applied to treat recurrent *Clostridioides difficile* infections (rCDI) possibly through bacteriophage-mediated (bacterial viruses, in short phages) modulation of the GM landscape [1–4]. The fecal donor material used for FMT is screened for pathogenic bacteria and viruses prior FMT to ensure the safety. However, this process is laborious and may only end up with 3% of the donor candidates passing all safety steps [5], and screening procedures fail, there is risk of transferring disease-causing agents as emphasised by a an incident June 2019 where two patients in the US had severe infections following FMT, of which one patient died [6]. As an alternative to FMT, fecal virome transplantation (FVT, sterile filtrated donor feces) has also shown promising efficacy against CDI and rCDI [2, 7, 8]. An important advantage of FVT over FMT is the elimination of bacterial transfer as a potential threat. However, there is still the risk of transferring disease-causing eukaryotic viruses despite the screening of donor material for known pathogenic viruses as long-term effects of most of the viruses inhabiting the human gastrointestinal tract are not yet studied [9, 10].

The interactions between gut bacteria and phages are complex and mutual, hence making the gut virome an important component in health and disease [11, 12]. Microbiome abundances and diversity are predictive of virome richness and diversity [13]. The estimated number of virus-like particles (VLP) remains between 10^9^ and 10^10^ VLP per gram of feces [14–17]. The human gut virome is dominated by phages (over 97%) while only about one tenth of those have been annotated until now [18]. The healthy adult gut virome is highly personal and stable over time within each individual [14, 19, 20]. Phages belonging to the order *Caudovirales* are together with single stranded DNA phages (ssDNA) of the order *Petitvirales* dominant in the human gut [10–12, 21, 22]. There are several indications that viromes of the dysbiotic and healthy gut microbiota differ [23–25], however the causal links between gut virome dysbiosis and disease is still poorly understood. However, the impact of the phageome on the composition and function of the gut microbiota has been suggested to have important consequences for health and outcome of FMT or FVT treatments [2, 26–29].

While phages only infect bacterial and not eukaryotic cells, we aimed to develop methodology to produce active enteric phage communities with minimized amounts of eukaryotic viruses using two different inocula; murine cecal and human fecal microbiota. Of which the chemostat propagated murine gut virome ability to improve gut microbiota balance was evaluated in preclinical studies representing animal models with distinct disease etiologies, CDI [8] and diet-induced obesity [30]. Our recent studies highlighted the importance of dilution rate on the composition of cultured fecal microbiota [31, 32]. The cultivation conditions and substrates were chosen to sustain the highest microbial diversity based on previous knowledge. Dynamics of the chemostat cultures was followed by determination of bacterial and viral composition, growth characteristics and metabolic products. We hypothesized that eukaryotic viruses would be washed out in chemostat culture due to the dilution effect. To our knowledge, the dynamic changes of enteric phages in chemostat cultures have not been described earlier.

## METHODS

### Chemostat inocula

Chemostat cultivations were carried out with two different intestinal inocula of mouse and human origin, respectively. Cecal contents of mice from different vendors were pooled, as previously we have shown that mice from different vendors represents distinctly different gut microbiota profiles (both the bacterial and viral community) [33, 34]. In total 18 C57BL/6N male mice were purchased to harvest intestinal content for downstream applications. The mice were five weeks old at arrival and purchased from three vendors, represented by 6 C57BL/6NTac mice (Taconic, Lille Skensved, Denmark), 6 C57BL/6NRj mice (Janvier, Le Genest-Saint Isle, France), and 6 C57BL/6NCrl mice (Charles River, Sulzfeld, Germany) and ear marked at arrival. Animal housing was carried out at Section of Experimental Animal Models, University of Copenhagen, Denmark, under the conditions as described previously [33]. For 13 weeks the mice were fed *ad libitum* low-fat diet (LF, Research Diets D12450J, USA) until termination at 18 weeks old and their body weight were measured every second week. To preserve the viability of the strict anaerobic bacteria, 6 mice from each vendor (in total 18 mice) were sacrificed by cervical dislocation and immediately transferred to a jar containing an anaerobic sachet (cat. no. AN0035A AnaeroGen, Thermo Fisher Scientific) and subsequently to an anaerobic chamber (containing ∼93 % N_2_, ∼2 % H_2_, ∼5 % CO_2_) at room temperature (Model AALC, Coy Laboratory Products, Grass Lake, Michigan, USA) where cecum content of the mice was sampled. Inside the anaerobic chamber, the samples were processed according to vendor (Janvier, Charles River and Taconic); weighted, suspended in an anoxic 1:1 mixture of PBS (NaCl 137 mM, KCl 2.7 mM, Na2HPO4 10 mM, KH2PO4 1.8 mM) and 50% glycerol and homogenized in BagPage® 100 mL filter bags (Interscience, Saint-Nom-la-Bretèche, France) with a laboratory stomacher (Stomacher 80, Seward, UK) at medium speed for 120 seconds. The cecum content from mice from all vendors were mixed, and the pooled cecum content was divided into 6 cryotubes ∼ 0.5 g cecum content in each, one for each chemostat run. The samples were frozen and kept at -80°C until use in chemostat experiments. The abovementioned processes are illustrated with a flow-diagram (Supplementary Fig. S1). All procedures regarding the handling of these animals used for donor materiel were carried out in accordance with the Directive 2010/63/EU and the Danish Animal Experimentation Act with the license ID: 2012-15-2934-00256.

The human study was approved by Tallinn Medical Research Ethics Committee, Estonia (protocol no. 554). Fecal samples from seven healthy donors (age 19-37 years, Caucasian, three male and four female) were diluted five times in dimethyl sulfoxide phosphate saline buffer, pooled in equal volumes and stored frozen at -80°C until use as described previously in Adamberg et al. [35].

### Growth medium

The base medium was prepared in 0.05 M potassium phosphate buffer containing amino acids, mineral salts and vitamins as described previously [32]. Hemin (5 mg/L), menadione (0.5 mg/L), bile salts (0.5 g/L), NaHCO3 (2.0 g/L), Tween-80 (0.5 g/L), Na-thioglycolate (0.5 g/L) and Cys-HCl (0.5 g/L, freshly made in oxygen reduced water) were added to the base medium. Carbohydrate sources and other components added to the medium for murine cultures were apple pectin (2 g/L, Sigma-Aldrich, USA), chicory inulin HP (1 g/L, Orafti, Belgium), corn core xylan (2 g/L, TCI, Japan), corn starch (5 g/L, Sigma-Aldrich, USA), larch wood arabinogalactan (2 g/L, TCI, Japan) and porcine mucin (4 g/L, Type II, Sigma Aldrich, USA), acetic acid (0.3 g/L, Sigma-Aldrich, USA), tryptone (3 g/L, LABM, UK) and yeast extract (3 g/L, LABM, UK) as described by Macfarlane et al. [36]. Carbohydrate sources for the human fecal matter inoculated cultures were apple pectin (2.5 g/L, Sigma-Aldrich, USA) and porcine mucin (2.5 g/L, Type II, Sigma Aldrich, USA). The carbohydrate sources were sterilized separately and added to the medium before experiments. The medium for mouse cecal matter inoculated cultures contained about three times more carbohydrates than that for human fecal matter inoculated cultures.

### Cultivation system and culture conditions

The cultivation system described earlier [32] was used for human fecal and mouse cecal matter inoculated cultures. Briefly, the Biobundle cultivation system consisting of fermenter, the ADI 1030 bio-controller and cultivation control program “BioXpert” (Applikon, The Netherlands) was used. The fermenter was equipped with sensors for pH, pO_2_, and temperature control. Variable speed pumps for feeding and outflow were controlled by a chemostat algorithm: D = F/V, where D is the dilution rate (1/h), F is the feeding rate (L/h), and V is the fermenter working volume (L). pH was controlled by adding 1M NaOH according to the pH setpoint. The medium in the feeding bottle and the culture were flushed with sterile-filtered nitrogen gas (99.9%, AS Linde Gas, Estonia) before inoculation and throughout the cultivation to maintain anaerobiosis. The culture volume was kept constant (600 ml for mouse cecal and 300 ml for human fecal matter inoculated cultures). The temperature was kept at 36.6 °C. pH was kept constant at pH=6.4 for mouse cecal and 7.0 for human fecal matter inoculated cultures depending on the physiological pH of the host. The scheme of experiments with mouse cecal matter inoculated culture is depicted in the Fig. 1. Three ml of the pooled mouse cecum matter were inoculated into 600 ml medium to start the experiments. The chemostat algorithm was started 15-20 hours after inoculation, which corresponds to the middle of the exponential growth phase of the fecal culture. Three replicates were carried out with human fecal and mouse cecal inocula at two dilution rates, 0.05 1/h (D_low_) and 0.2 1/h (D_high_), except for experiments with mouse cecal inocula at D_high_ where two experiments were performed. The stabilization of five residence times was used in all experiments. On-line and at-line parameters used for experiment control are depicted on the Supplementary Fig. S2.

**Fig. 1.**
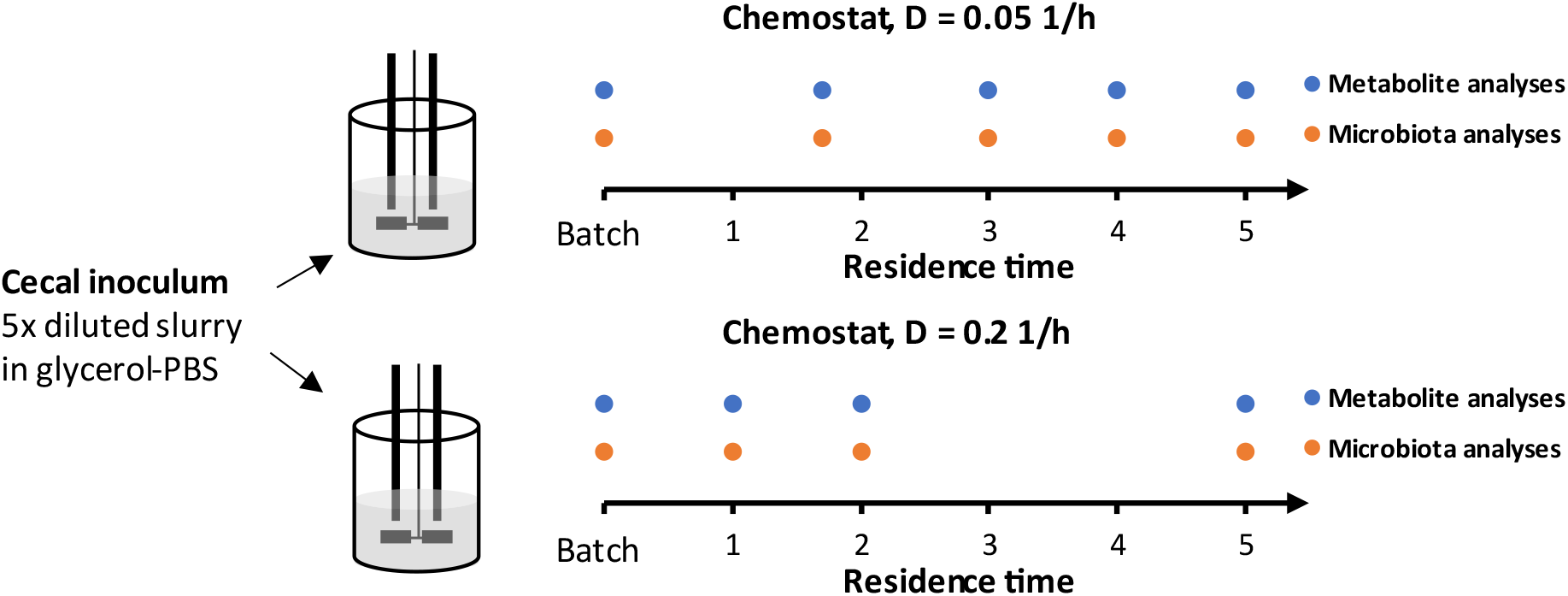
The experimental setup of chemostat cultivations of mice cecal culture. At start one percent of five times diluted cecal content was inoculated into a bioreactor followed by batch growth 23 h. The chemostat culture was then run up to 5 residential times. Colored dots indicate the sampling points for metabolite and microbiota (16S rRNA gene amplicon and virome sequencing) analyses. Similar setup was used for human fecal cultures [32].

### Analytical methods

Samples from the outflow were collected on ice, centrifuged (14,000 g, 5 min, 4 °C) and stored separately as pellets (at -80°C) and supernatants (at -20°C) until further analyses. For chromatographic analyses, culture supernatants were filtered using AmiconR Ultra-10K Centrifugal Filter Devices, cut-off 3 kDa according to the manufacturer’s instructions (Millipore, USA). The concentrations of organic acids (succinate, lactate, formate, acetate, propionate, isobutyrate, butyrate, isovalerate and valerate) and ethanol were determined by high-performance liquid chromatography (HPLC, Alliance 2795 system, Waters, Milford, MA, USA), using BioRad HPX-87H column (Hercules, CA, USA) with isocratic elution of 0.005 M H2SO4 at a flow rate of 0.5 mL/min and at 35 °C. Refractive index (RI) (model 2414; Waters, USA) and UV (210 nm; model 2487; Waters, USA) detectors. Analytical grade standards were used for quantification of the substances. The detection limit for the method was 0.1 mM.

The composition of the gas outflow (H_2_, CO_2_, H_2_S, CH_4_, and N_2_) was analyzed using an Agilent 490 Micro GC Biogas Analyzer (Agilent 269 Technologies Ltd., USA) connected to a thermal conductivity detector. The volume of the gas flow was regularly recorded using MilliGascounter (RITTER Apparatebau GMBH & Co, Germany).

The Redox potential of the growth medium and culture supernatant was measured by a pH/Redox meter using an InLab®Redox electrode (Mettler Toledo, USA). The biomass dry weight was measured gravimetrically from 10 ml culture by centrifugation (6,000 rpm, 20 min), washing the biomass with distilled water and drying in an oven at 105°C for 20 h.

### Pre-processing of samples for separation of viruses and bacteria

Culture and inoculum samples were included to investigate microbiome changes over time. Separation of the viruses and bacteria from the culture/inoculum samples generated a pellet and supernatant by centrifugation and 0.45 µm filtering as described previously [33]. The volume of culture/inoculum homogenate was adjusted to 5 mL using SM buffer.

### Bacterial DNA extraction, sequencing and pre-processing of raw data

Human gut bacteriome data have been previously published [32] but to minimize errors due to using different data processing methods, human gut bacteriome sequences were reanalyzed using the same pipeline as described above for analysis of mice samples. Bead-Beat Micro AX Gravity (mod.1) kit from A&A Biotechnology (Gdynia, Poland) was used to extract bacterial DNA from the culture/fecal pellet by following the instructions of the manufacturer. The final purified DNA was stored at -80<u>°</u>C and the DNA concentration was determined using Qubit HS Assay Kit (Invitrogen, Carlsbad, California, USA) on the Qubit 4 Fluorometric Quantification device (Invitrogen, Carlsbad, California, USA). The bacterial community composition was determined by NextSeq-based (Illumina) high-throughput sequencing (HTS) of the 16S rRNA gene V3-region, as previously described [24]. Quality-control of reads, de-replicating, purging from chimeric reads and constructing zOTU was conducted with the UNOISE pipeline [37] and taxonomically assigned with Sintax [38]. Taxonomical assignments were obtained using the EZtaxon for 16S rRNA gene database [39]. Code describing this pipeline can be accessed in github.com/jcame/Fastq_2_zOTUtable. The average sequencing depth after quality control (Accession: PRJEB58777, available at ENA) for the fecal 16S rRNA gene amplicons was 60,719 reads (min. 11,961 reads and max. 198,197 reads).

### Viral RNA/DNA extraction, sequencing and pre-processing of raw data

The sterile filtered culture/fecal supernatant was concentrated using centrifugal filters Centrisart with a filter cut-off at 100 kDA (Sartorius) by centrifugation centrifuged at 1,500 x g at 4<u>°</u>C (dx.doi.org/10.17504/protocols.io.b2qaqdse). The viral DNA/RNA was extracted from the culture/fecal supernatants using the Viral RNA mini kit (Qiagen) as previously described [33]. Reverse transcription was executed using the SuperScript VILO Master mix by following the instructions of the manufacturer and subsequently cleaned with DNeasy blood and tissue kit (Qiagen) by only following step 3-8 of the manufacturers standard protocol. In brief, the DNA/cDNA samples were mixed with ethanol, bound to the silica filter, washed two times, and eluted with 40 µL elution buffer. Multiple displacement amplification (MDA, to include ssDNA viruses) using GenomiPhi V3 DNA amplification kit (Cytiva) and sequencing library preparation using Nextera XT kit was performed at previously described [33], and sequenced at a commercial facility using the NovaSeq platform (NovoGene). The average sequencing depth of raw reads (Accession: PRJEB58787, available at ENA) for the fecal viral metagenome was 22,701,135 reads (min. 342,022 reads and max. 203,403,294 reads. The raw reads were trimmed from adaptors and the high-quality sequences (>95% quality) using Trimmomatic v0.35 [40] with a minimum size of 50 nt were retained for further analysis. High quality reads were de-replicated and checked for the presence of PhiX control using BBMap (bbduk.sh) (https://www.osti.gov/servlets/purl/1241166). Virus-like particle-derived DNA sequences were subjected to within-sample de-novo assembly-only using Spades v3.13.1 [41] and the contigs with a minimum length of 2,200 nt, were retained. Contigs generated from all samples were pooled and de-replicated at 90% identity using BBMap (dedupe.sh). Prediction of viral contigs/genomes was carried out using VirSorter2 [42] (“full” categories | dsDNAphage, ssDNA, RNA, Lavidaviridae, NCLDV | viralquality ≥ 0.66), vibrant [43] (High-quality | Complete), and checkv [44] (High-quality | Complete). Taxonomy was inferred by blasting viral ORF against viral orthologous groups (https://vogdb.org) and the Lowest Common Ancestor (LCA) for every contig was estimated based on a minimum e-value of 10e^-5^. Phage-host prediction was determined by blasting (85% identity) CRISPR spacers and tRNAs predicted from >150,000 gut species-level genome bins (SGBs) [45, 46]. Following assembly, quality control, and annotations, reads from all samples were mapped against the viral (high-quality) contigs (vOTUs) using the bowtie2 [47] and a contingency-table of reads per Kbp of contig sequence per million reads sample (RPKM) was generated, here defined as vOTU-table (viral contigs). Code describing this pipeline can be accessed in github.com/jcame/virome_analysis-FOOD. Viral mock communities (phage C2, T7, P35, MS2, Phi6, PhiX174, T4, and PMBT5) were used as positive controls to evaluate if the library preparation and sequencing could detect ss/dsDNA and ss/dsRNA viruses with a titer of ∼10^6^ PFU/mL from both pure phage culture as well as spiked fecal matrices.

### Bioinformatic analysis of bacterial and viral sequences

Initially the dataset was purged for zOTU’s/viral contigs, which were detected in less than 5% of the samples, but the resulting dataset still maintained 99.5% of the total reads. Cumulative sum scaling (CSS) [48] was applied for the analysis of beta-diversity to counteract that a few zOTU/viral contigs represented a large fraction of count values, since CSS have been benchmarked with a high accuracy for the applied metrics [49]. CSS normalization was performed using the R software using the metagenomeSeq package [50]. R version 4.2.2 [51] was used for subsequent analysis and presentation of data. The main packages used were phyloseq [52], vegan [53], deseseq 2 [54], ampvis2 [55], ggpubr, and ggplot2 [56]. A-diversity analysis was based on raw read counts and statistics were based on ANOVA. B-diversity was represented by Bray Curtis dissimilarity metrics and statistics were based on PERMANOVA.

## RESULTS

With the aim of generating viromes depleted of eukaryotic viruses, mouse cecal and human fecal matter were propagated initially as batch cultures and subsequently in chemostat mode at two different dilution rates (D_low_, 0.05 1/h and D_high_, 0.2 1/h) for five residence times. The overall concept for both cultures (i.e dilution rates, temperature, and pH) was the same except for some differences in the media composition. Most importantly, the total carbohydrate concentration was three times higher in the medium for mouse-mimicking conditions compared to that for the human conditions mimicking cultures (15.2 g/L vs 5 g/L, respectively), to resemble the high content of complex carbohydrate in mice chow feed.

### Wash-out of eukaryotic viruses from chemostat cultures

The relative abundance of eukaryotic viruses decreased already after batch phase and were almost depleted after five residence times in chemostats of both mouse cecal and human fecal matter inoculated cultures (Fig. 2). The relative abundance of eukaryotic viruses after batch phase was 0.3±0.2 % and 0.4±0.2 % (average ± standard deviation) in mouse cecal and in human fecal matter inoculated cultures, respectively. During the chemostats relative abundance of eukaryotic viruses declined to 0.006 % and 0.04 % in mice cecal and human fecal matter inoculated cultures at slow dilution rates, respectively. At the same time, the bacterial virome maintained high diversity in the chemostat phase and the number of bacterial viral OTUs remained at least 1000 times higher than these of eukaryotic viral OTUs (Fig. 3 and Supplementary Fig. S3).

**Fig. 2.**
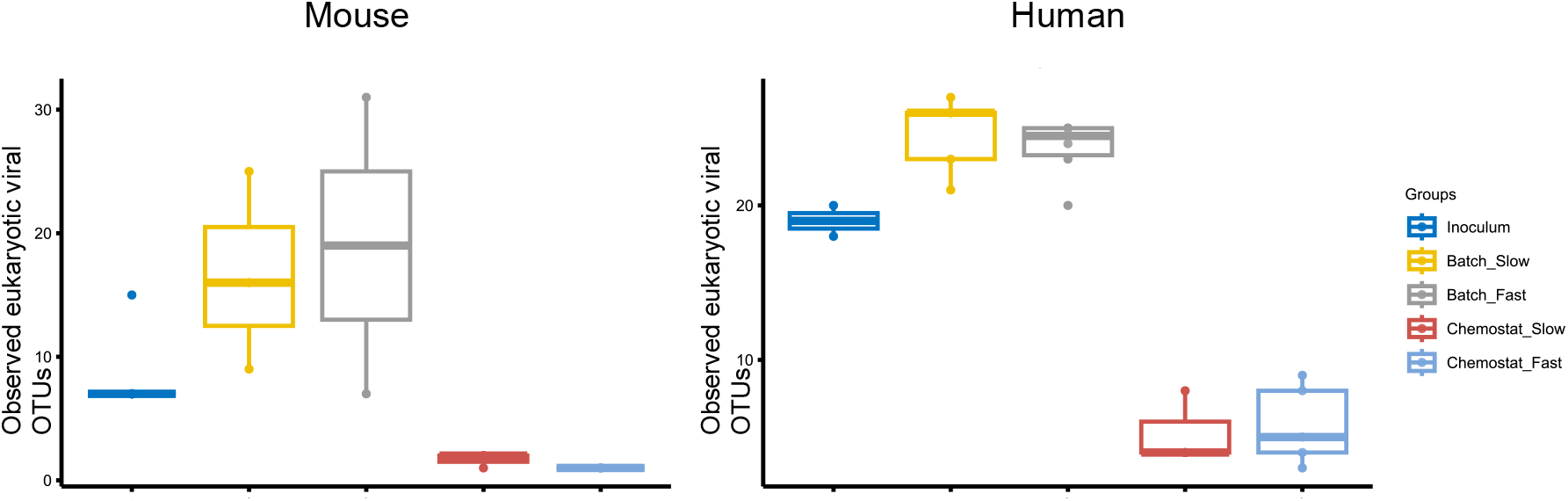
Chemostat propagation of cecal/ fecal inocula leads to depletion of eukaryotic viruses shown as the number or observed eukaryotic viral operational taxonomic units (vOTUs). Batch_Slow and Batch_Fast designate the batch cultures prior the chemostat mode of low and fast dilution rates, respectively.

**Fig. 3.**
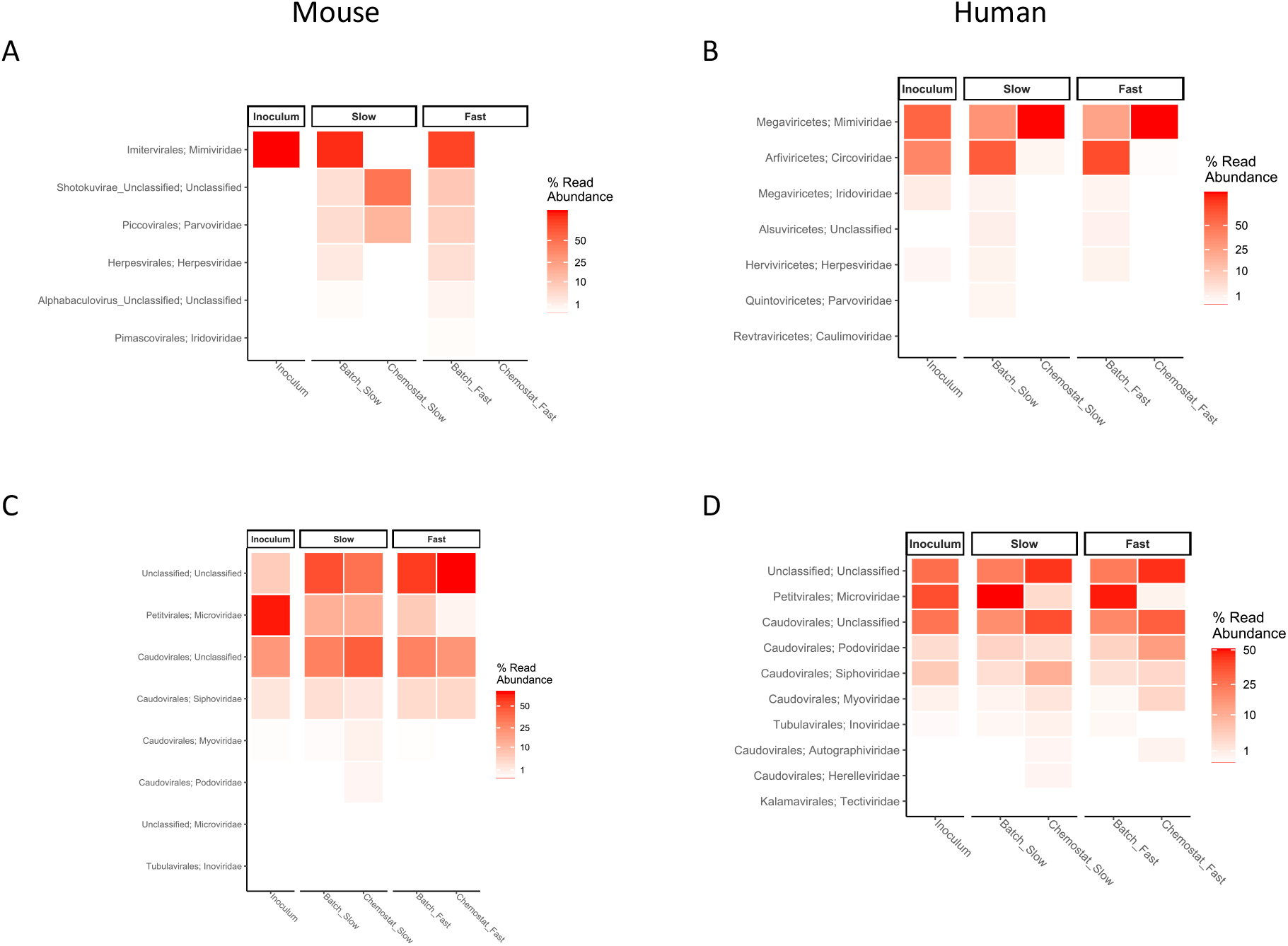
Diversity of prokaryotic viruses during chemostat propagation of mouse cecal/ human fecal matter inoculated cultures. Depletion of eukaryotic viral families and enrichment of bacterial viral families from batch to stabilized chemostat cultures (A and C – mouse cecal matter inoculated cultures, B and D – human fecal matter inoculated cultures) presented as heatmaps. Color intensity indicates the relative abundance of eukaryotic (A, B) or prokaryotic (C, D) viral contigs from total eukaryotic or prokaryotic viral contigs, respectively. ‘Slow’ and ‘Fast’ indicate the dilution rates used in the chemostat (D_low_ 0.05 and D_high_ 0.2 1/h, respectively). Batch_Slow and Batch_Fast designate the batch cultures prior the corresponding chemostat modes of low and fast dilution rates, respectively.

Both eukaryotic DNA viruses and phages were identified in both inocula (Fig. 3). RNA viruses (*Leviviridae* and *Cystoviridae*) were found only in human feces and their count in mouse cecal matter remained below detection limit of virus particles (10^6^ VLP/mL). The most abundant taxon of eukaryotic viruses in mouse virome was *Mimiviridae* and the abundances of other identified viral taxa (*Herpesviridae* and *Parvoviridae*) were two magnitudes lower than *Mimiviridae*. Similarly, *Mimiviridae* and *Circoviridae* were the most abundant eukaryotic viruses in human feces. However, their relative abundance from all viruses was below 1%. Of note, nearly half of the vOTUs remained unidentified in both mouse and human inocula.

### Reproducibility of the chemostat-cultured viromes

Chemostat experiments showed that bacterial and archaeal viruses were persistent to continuous culturing of gut microbiota and showed the dilution rate specific patterns. The diversity of the bacterial virome was significantly related to the dilution rate (Fig. 4). The Shannon index of viromes of mouse cecal matter inoculated cultures remained high even at the end of low dilution rate chemostat while remarkable reductions of diversity were observed at high dilution rate. In cultures inoculated with fecal matter of human origin, no clear associations were seen between the virome diversity and cultivation conditions. However, Shannon indices of bacteriomes of human fecal matter inoculated cultures were higher in chemostats than these in batch cultures (Supplementary Fig. S4). The main archaeal viral taxon was *Bicaudavirus* and its abundance was relatively stable in batch and chemostat modes in both mouse cecal and human fecal matter inoculated cultures (data not shown).

**Fig. 4.**
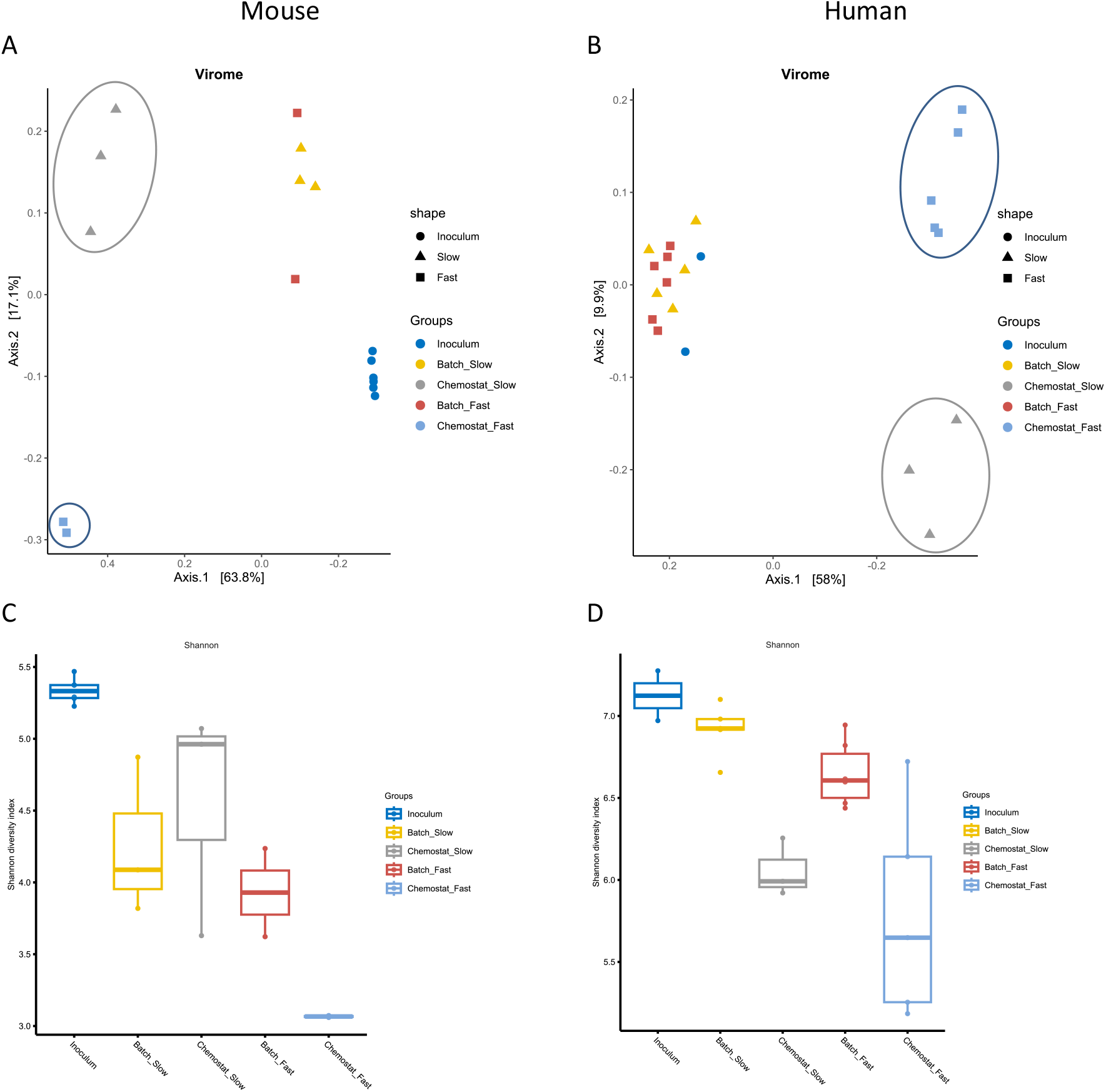
Chemostat propagation of fecal inocula at different dilution rates leads to viromes with reproducible composition shown by beta diversity (A, B) and diverse viral community shown by Shannon index (C, D). Samples from inoculum, batch and chemostat cultures of mouse cecal (A, C) and human fecal (B, D) matter inoculated cultures are shown. ‘Slow’ and ‘fast’ in the column names indicate the dilution rate used in the chemostat (D_low_ 0.05 and D_high_ 0.2 1/h, respectively). Batch_Slow and Batch_Fast designate the batch cultures prior the corresponding chemostat modes of low and fast dilution rates, respectively. Beta diversity and Shannon indices of bacteriomes are shown on the Supplementary Fig. S4.

When examining overall virome and bacteriome compositional patterns, Fig. 4 shows that inocula and samples of batch, slow and fast chemostat propagation clustered into distinct groups. Viromes of batch samples were closer to these of inoculum samples as there were no outflow but also show persistence of eukaryotic viruses in these conditions. More importantly, our results indicate good reproducibility of chemostat experiments as clearly separated clusters of chemostat samples were observed. Similar distinct clustering was observed also for the bacteriome (see below and Supplementary Fig. S4).

### The effect of dilution rate on the composition of bacteriomes and corresponding viromes

In mouse cecal matter inoculated batch cultures more than 90% of viral taxa could be identified while in chemostats the number of unidentified viruses increased up to 50% at both dilution rates. In the end of the batch phase, the bacteriome of the mouse cecal matter inoculated culture was dominated by *Bacteroides*, lactobacilli and *Enterobacteriaceae*. Correspondingly, the main taxa of viruses were *Microviridae* and *Siphoviridae* (Fig. 3), which include many viruses related to *Escherichia* and lactobacilli [57, 58]. *Microviridae*, the most abundant taxa in the inoculum decreased significantly already in the batch phase and were mostly washed out for the end of fast chemostat. However, in slow chemostat, *Microviridae* were still present in noticeable amounts by the end of experiment. Other prevalent taxa in the virome at low dilution rate were *Myoviridae* and *Podoviridae*.

The most abundant vira in human fecal matter inoculated cultures were also *Microviridae* (more than 70%) followed by *Siphoviridae* and *Podoviridae* while the fraction of unidentified viruses remained below 10%. In the end of batch phase the relative abundance of *Microviridae* and other annotated phages decreased while that of unidentified viruses increased several times compared to their proportions in the inocula. At both dilution rates of chemostat cultures, *Microviridae* were mostly washed out and largely replaced by viruses from the family *Myoviridae, Podoviridae*, and *Siphoviridae. Myoviridae* and *Podoviridae* were especially abundant at high dilution rate.

The assembled viral contigs were used to predict bacterial hosts using databases of CRISPR-arrays and tRNA profiles. According to 16S rRNA gene sequence data, the dominant taxa in the mouse inoculum were *Ruminococcaceae, Lachnospiraceae* and lactobacilli, which by the end of the batch phase had changed to *Bacteroides, Lactobacillus, Escherichia* and *Enterobacter* (Fig. 5). In the chemostat cultures with high dilution rates (D = 0.2 1/h), the abundance of these four genera remained high. The abundance of lactobacilli decreased from 23% to 12% while that of *Bacteroides* and enterobacteria remained the most dominant taxa (both more than 20 %) in the end of chemostat propagation. In chemostats with low dilution rate inoculated with mouse cecal matter, *Bifidobacterium, Bacteroides, Blautia* and an unidentified *Ruminococcaceae* became dominant. At the same time lactobacilli and *Escherichia* were washed out while *Akkermansia, Intestinimonas* and an unidentified *Lachnospiraceae* took over their place. Based on the virome host prediction analyses, the prevalence of *Bacteroides* related viruses decreased more than five times at low dilution rate being negligible at the end of chemostat propagation (Supplementary Fig. S4). Higher bacteriome diversity was reflected also in higher virome diversity. There were viruses related to genera *Akkermansia, Blautia, Enterococcus* and *Lachnoclostridium*, but viruses related to bifidobacteria and *Ruminococcaceae* were not detected (Supplementary Fig. S4).

**Fig. 5.**
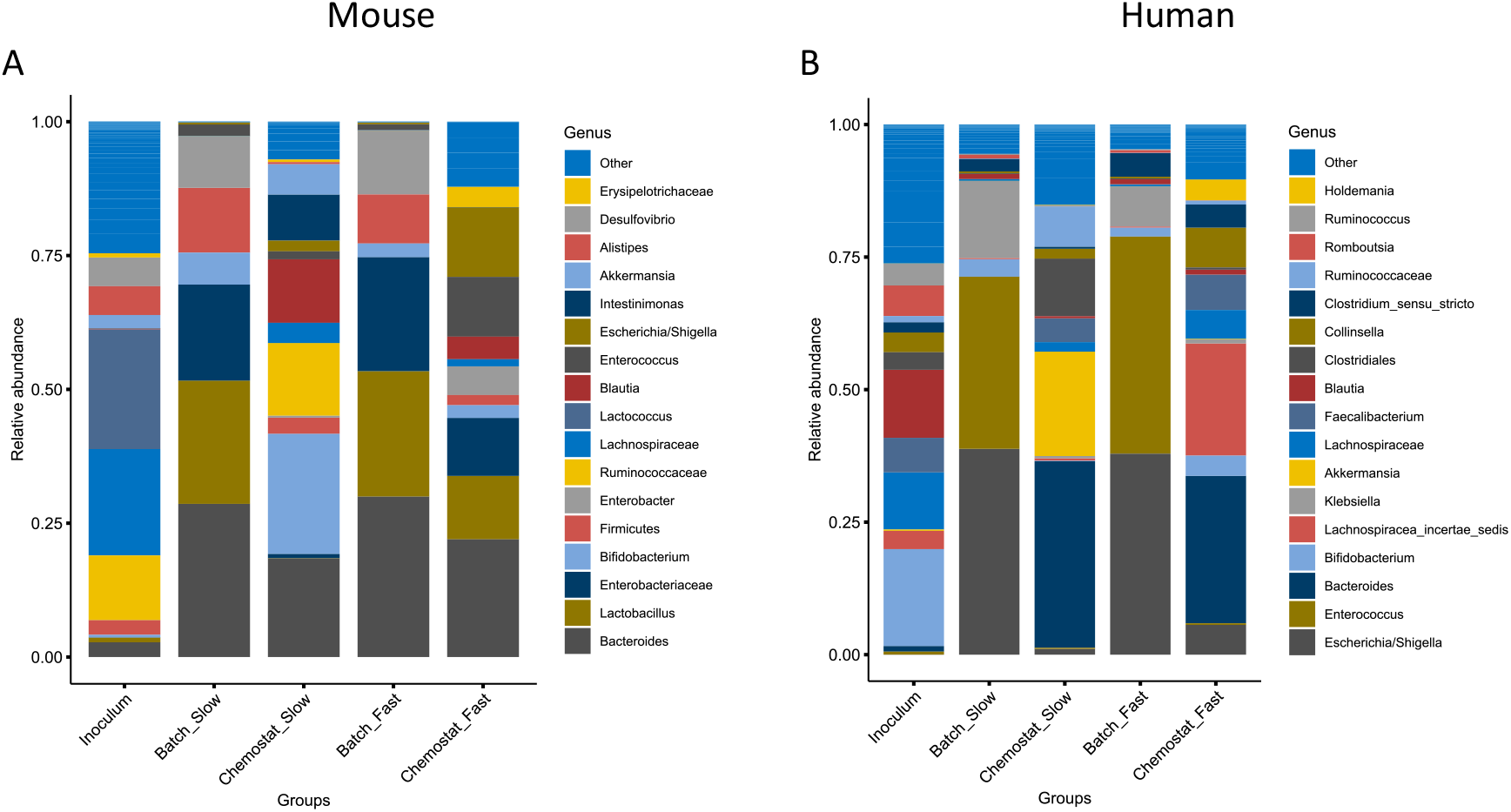
Bacteriome composition of mice cecal (A) and human fecal matter inoculated cultures (B) after batch and chemostat propagation. ‘Slow’ and ‘fast’ in the column names indicate the dilution rate used in the chemostat (D_low_ 0.05 and D_high_ 0.2 1/h, respectively). Batch_Slow and Batch_Fast designate the batch cultures prior the corresponding chemostat modes of low and fast dilution rates, respectively.

Completely different patterns could be seen in chemostats inoculated with fecal matter of human origin. These were dominated by *Escherichia* and *Enterococcus* in the end of batch phase. It can be explained by the remarkable differences of the composition of human fecal and mouse cecal inocula although the dominant bacterial taxa were comparable. In steady state (end of chemostat), the composition of cultured mouse and human microbiota were more similar. *Bacteroides*, enterobacteria, bifidobacteria and *Clostridium* were abundant in both cultures at high dilution rate while dominance of several *Lachnospiraceae, Faecalibacterium* and *Collinsella* was characteristic to human fecal matter inoculated cultures only. On the other hand, *Akkermansia* and *Ruminococcaceae* members were always characteristic of low dilution rate chemostats inoculated with either mouse cecal or human fecal matter. Virome host analysis showed that human fecal matter inoculated cultures contained mainly viruses related to *Bacteroides, Faecalibacterium* and *Prevotella* in the end of batch phase (Supplementary Fig. S4). The fast dilution rate promoted propagation of viruses related to *Bacteroides* but also to *Parabacteroides* and *Methanobrevibacter*. The latter was not detected in bacteriome analyses of 16S rRNA gene amplicon sequences.

### Bacterial metabolites were in accordance with the composition of microbiota

In mouse cecal matter inoculated batch cultures, the major metabolic products were acetate and lactate, comprising about 30 and 24 mol%, respectively, followed by formate, ethanol, succinate and propionate (Fig. 6). Butyrate was almost missing, and propionate represented less than 5 % of total acids. Such metabolite pattern is in accordance with the abundance of lactobacilli, *Bacteroides, Enterobacteriaceae* members, and *Bifidobacterium* in mouse cecal matter inoculated batch cultures (Fig. 5).

**Fig. 6.**
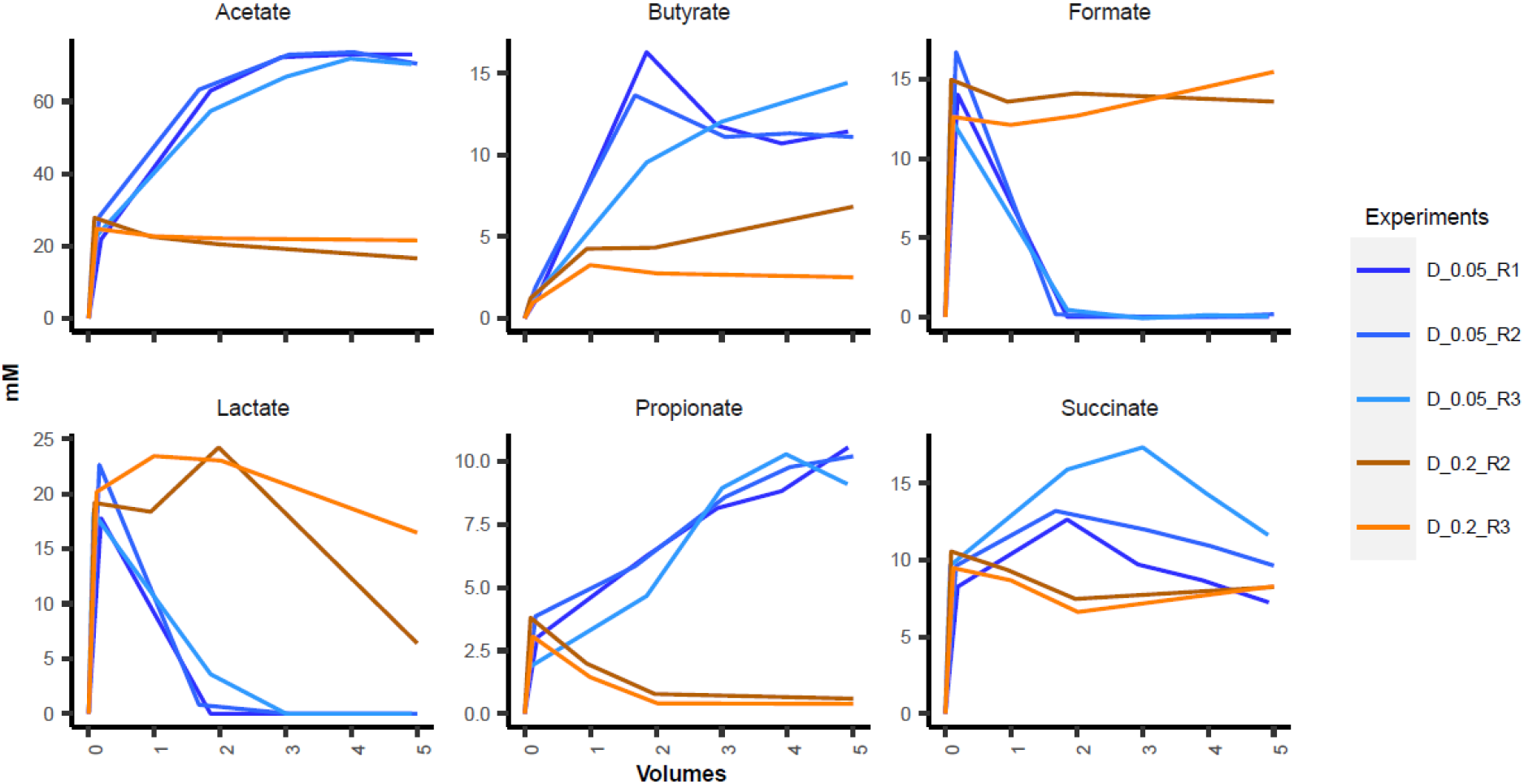
Reproducibility of metabolite dynamics of chemostat propagated mouse cecal cultures at two dilution rates. Metabolite concentrations are shown in millimoles in relation to residence time (‘Volumes’ on x-scale). Low dilution rate (D = 0.05 1/h) is indicated with blue lines and high dilution rate (D = 0.2 1/h) with brown-orange lines. R indicates the number of the replicate.

Remarkable differences in product profiles were observed of mouse cecal matter inoculated cultures at two dilution rates, D_low_ = 0.05 1/h and D_high_ = 0.2 1/h (Fig. 6). The highest acetate production (61 ± 2 mol-% of total products) was at D_low_, while lactic acid was not detected. Ethanol was produced in comparable amount with butyrate and propionate (all around 10 mol-%). Production of succinate was below 10 mol-% at D_low_ and the amount of formate was marginal (0.1 mol-%) while hydrogen sulfide was mainly detected at D_low_ (1.3 ± 0.1 mmol/gDW) compared to that at D_high_ (0.33 ± 0.03 mmol/gDW). Metabolite data on the human fecal matter inoculated cultures are presented in a recent paper by Adamberg et al [32]. When comparing the cultures of the mouse cecal and human fecal matter inoculated cultures, the overall acid and gas production patterns were similar at high dilution rate (D_high_) but differed at low dilution rate (D_low_). However, propionate production was higher at D_low_ and lactate production was minor at D_high_ in human fecal matter inoculated cultures [32].

## DISCUSSION

### Reproducibility of the virome propagation

Here we aimed to produce chemostat-propagated gut viromes depleted of eukaryotic viruses. Two different inocula (mouse cecal and human fecal matter) were grown at two different dilution rates, D_low_ = 0.05 and D_high_ = 0.2 1/h. The median relative abundance of contigs of eukaryotic origin decreased from 0.8 % and 0.6 % (mouse cecal and human fecal matter inoculated cultures, respectively) to less than detectable amounts after five residence times in chemostat cultivation. Interestingly, the viromes appeared to be reproducible in replicate runs as both the bacteriomes and the viromes approached equilibrium after five residence times (Supplementary Figs. S3 and S5). It would be expected that extended time of the chemostat process remove more eukaryotic viral particles while maintaining a similar phageome profile. This approach provides a safe and potential effective phage-mediated gut microbiota modulation tool to investigate or treat gut-associated diseases.

Very little is known concerning virus profiles of continuous cultures inoculated with fecal or cecal matter. However, the composition of the virome would be expected to be reflected by the bacteriome profile due to their inherent host-phage relationship. Interestingly, the relative abundances of phages did not correspond to their host bacteria in the bacteriome as seen from Figs. 3 and 5. In our study *Siphoviridae* was the dominant viral family in both mouse cecal and human fecal matter inoculated cultures, whilst *Podoviridae* and *Myoviridae* were abundantly present in the human fecal matter inoculated chemostat culture with high dilution rate. *Microviridae*, the most abundant viral family In both inocula was almost washed out in chemostat cultures, excluding slow chemostats started with mouse cecal matter inoculum. High abundance of *Microviridae* in the virome can be explained by method used for sample preparation. Multiple displacement amplification (MDA) has a preference for ssDNA viruses, which might inflate *Microviridae* numbers [59, 60], however, Shah et al. showed recently that using short MDA (half an hour) the data were still quantitative as confirmed by plaque assays of double-stranded DNA (dsDNA) *Escherichia coli* infecting viruses [10].

### Impact of dilution rate

Dilution rate is a critical factor that determines the composition and metabolism of the chemostat propagated bacteria [36]. This is in line with the human gut transit time (a sort of “dilution rate”) being highly correlated to the gut microbiota diversity and composition [61, 62]. Dilution rates 0.2 and 0.05 1/h denote the fast degradation of dietary fibers in the cecum and slow grow rate in the colon [31]. Microbial metabolism directs the cross feeding between different community members and its changes affects the overall metabolic patterns and composition of the consortium. As an example, *Akkermansia*, a mucus-associated bacterium in the colon [63] was a dominant species in both mouse cecal and human fecal matter inoculated cultures at low dilution rate (0.05 1/h). On the other side the relative abundance of butyric acid producers (species of *Lachnospiraceae*) was higher at high dilution rate (0.2 1/h) [32]. In slow growing mouse cecal matter inoculated cultures higher diversity of the bacteriome was obtained similar to what has also been reported for human fecal matter inoculated chemostats by us and Asnicar et al. [62]. Additionally, high diversity of viromes at D_low_ were shown in this study. Thus, D_low_ in other words appear appropriate to produce diverse microbiomes and phageomes from gut microbiota cultures. In contrast, the growth of *Enterobacteriaceae* and viruses predicted to have *Enterobacteriaceae* as host was remarkably supported by fast dilution (D_high_ = 0.2 1/h) in mouse cecal matter inoculated chemostat-fermentations, probably reflecting the high maximum specific growth rate of *Enterobactericeae* [64].

### Chemostat propagation of active virome for *in vivo* trials

We have shown that chemostat can be used to produce reproducible bacterial consortia from human fecal matter inoculated cultures [32]. In this study we confirmed this phenomenon on mouse cecal matter inoculated cultures and showed also that chemostat can be used to produce reproducible viromes. The chemostat approach showed promising results for propagation of virome with minimal content of eukaryotic viruses [8, 30]. Fecal microbiota transplantation (FMT) allows modification of the intestinal microbiota in medical practice. Although FMT has been shown safe and effective in an immunocompromised CDI cohort [65], for these patients FMT is associated with several threats due to transfer of virulent microbes and viruses [66]. To overcome these potential risks, continuous cultivation of fecal inoculum can be used to diminish the load of eukaryotic viruses by dilution. In the absence of eukaryotic hosts chemostat cultures allow the propagation of bacteria and phage communities. During continuous growth over five volumes the bacterial and viral communities reach stabilized compositions that depend on the cultivation conditions (e.g dilution rate, pH, substrates).

In conclusion, we showed that chemostat cultivation is a highly promising method to generate mouse cecal and human fecal phageomes with minimal content of eukaryotic viruses. The phage populations can be used in transplantation experiments after removing all bacteria. Using the conditions tested here, the number of eukaryotic viruses can be decreased by more than hundred times of the initial load. This proof-of-concept study may constitute the first step of developing therapeutic tools to target a broad spectrum of gut-related diseases and thereby replacing FMT with a safer phage-mediated therapy.

