## Supplementary Figures for "Reproducible chemostat cultures to eliminate eukaryotic viruses from fecal transplant material"

### Supplementary materials

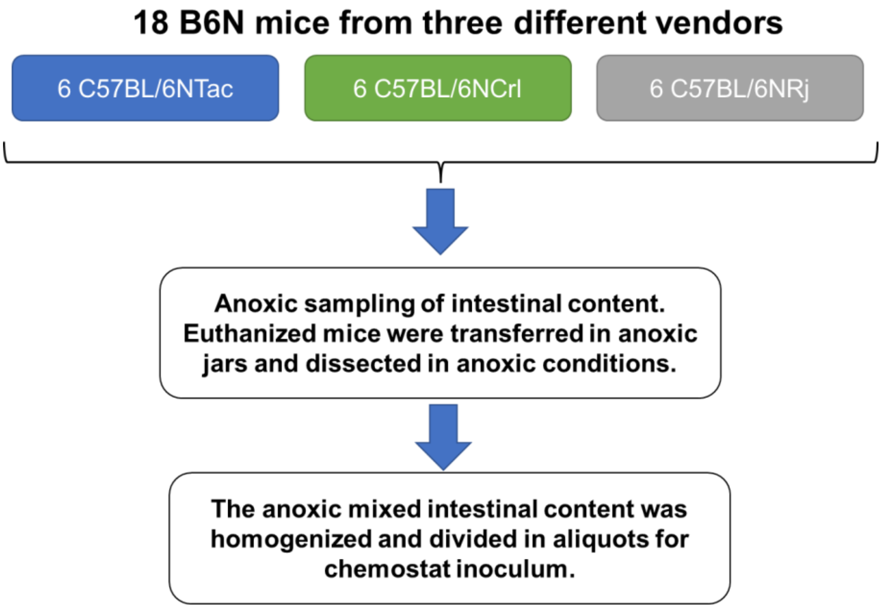


Supplementary Fig. S1. Flow diagram illustrating the origin of the intestinal donor content used for mouse inoculum. Initially, 18 mice from 3 different vendors (C57BL/6N-Tac, -Crl, -Rj) were sacrificed and their intestinal content from the cecum was collected in anoxic conditions to maintain the viability of the strict anaerobic bacterial GM members. The anoxic atmospheric conditions were maintained throughout the process. Regardless of vendor, the intestinal content was mixed and homogenized and divided into aliquots for chemostat inoculum.

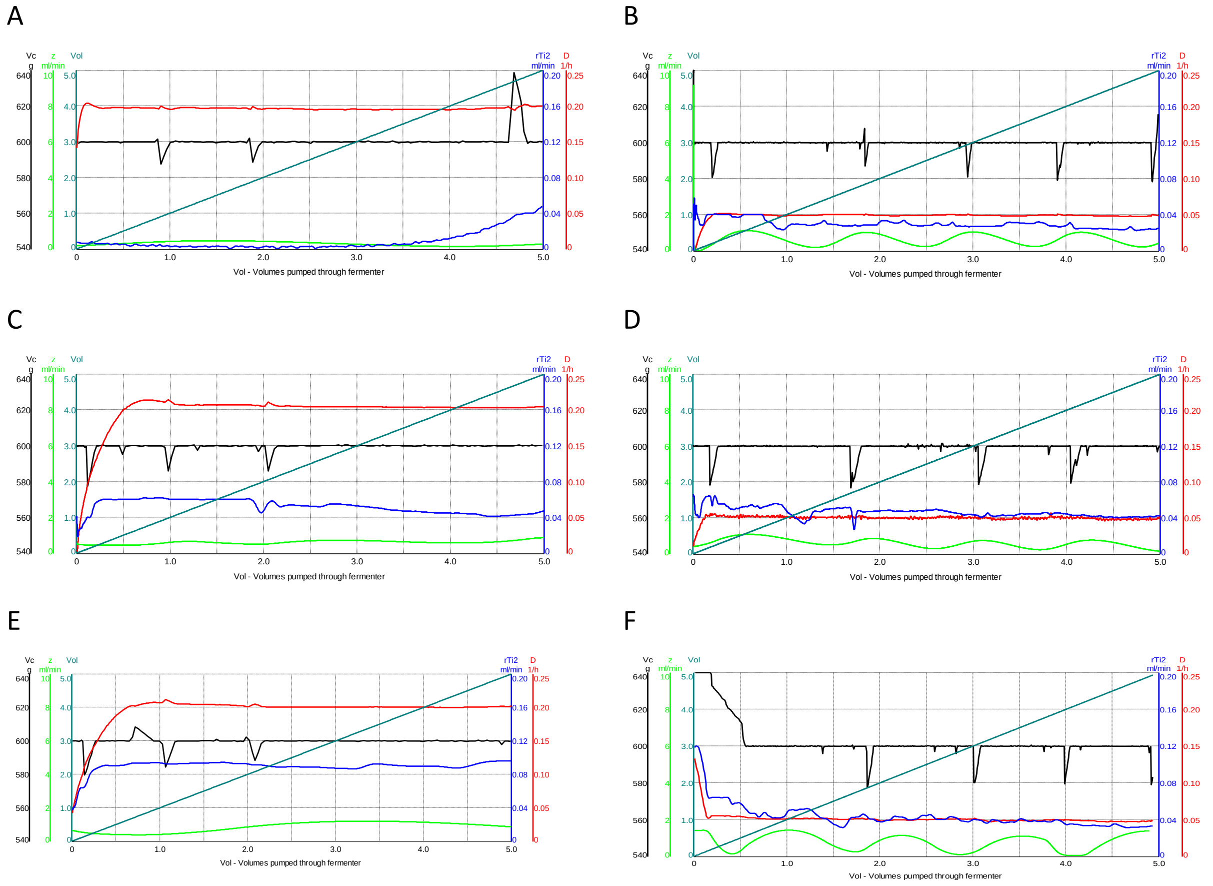


Supplementary Fig. S2. Dynamics of experimental parameters during chemostat cultivation of mouse cecal cultures. Figures A, C and E show data of three replicates at dilution rate 0.2 1/h and figures B, D and F at 0.05 1/h. rTi2 – titration rate of 1M NaOH (ml/min), Vc – fermenter volume, z – microbial gas production rate (ml/min) calculated as the difference of GasS-Gas, where GasS – total gas flow rate (N2+microbial), Gas – amount (ml) of nitrogen gas flown through the fermenter, D – dilution rate (1/h), Vol – indicates residence times.

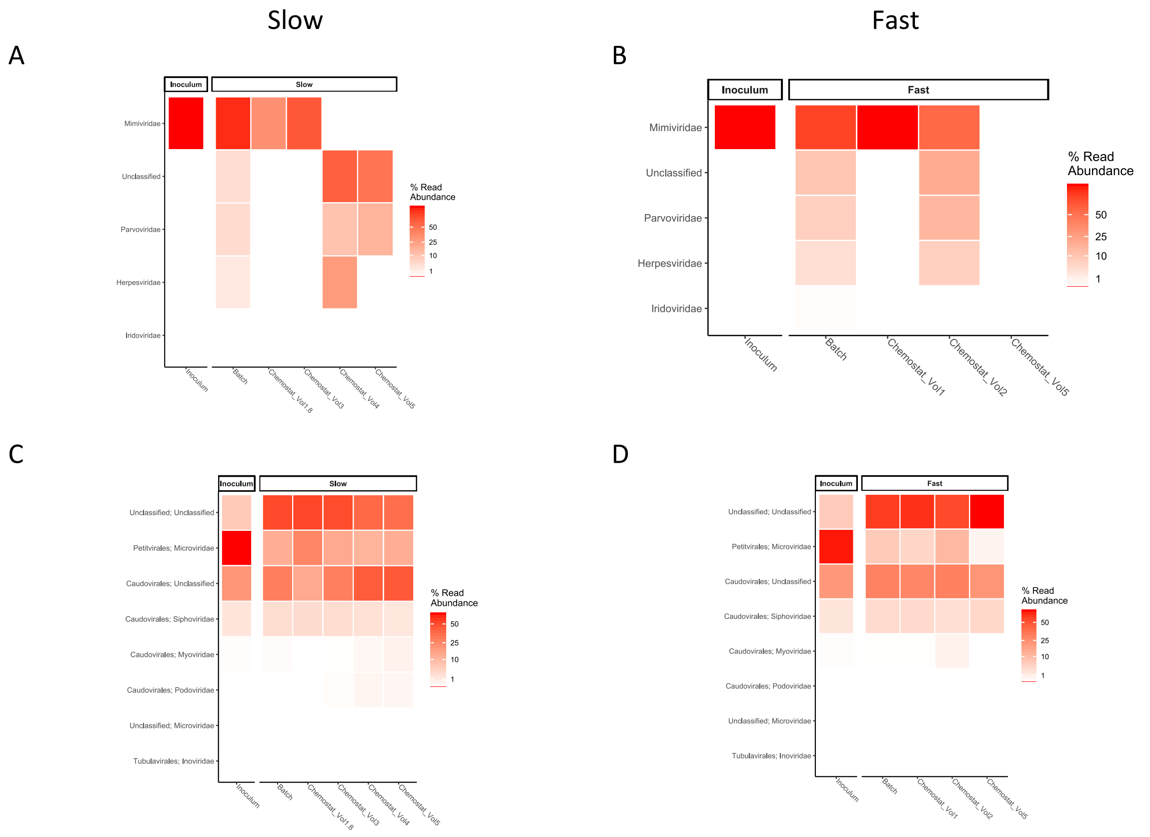


Supplementary Fig. S3. Stabilization of eukaryotic (A, C) and bacterial (B, D) viromes of mouse cecal cultures after switch from batch to chemostat mode. ‘Slow’ and ‘fast’ in the column names indicate the dilution rate used in the chemostat ($D_{low}$ 0.05 and $D_{high}$ 0.2 1/h, respectively). ‘Vol’ indicates the residence times.

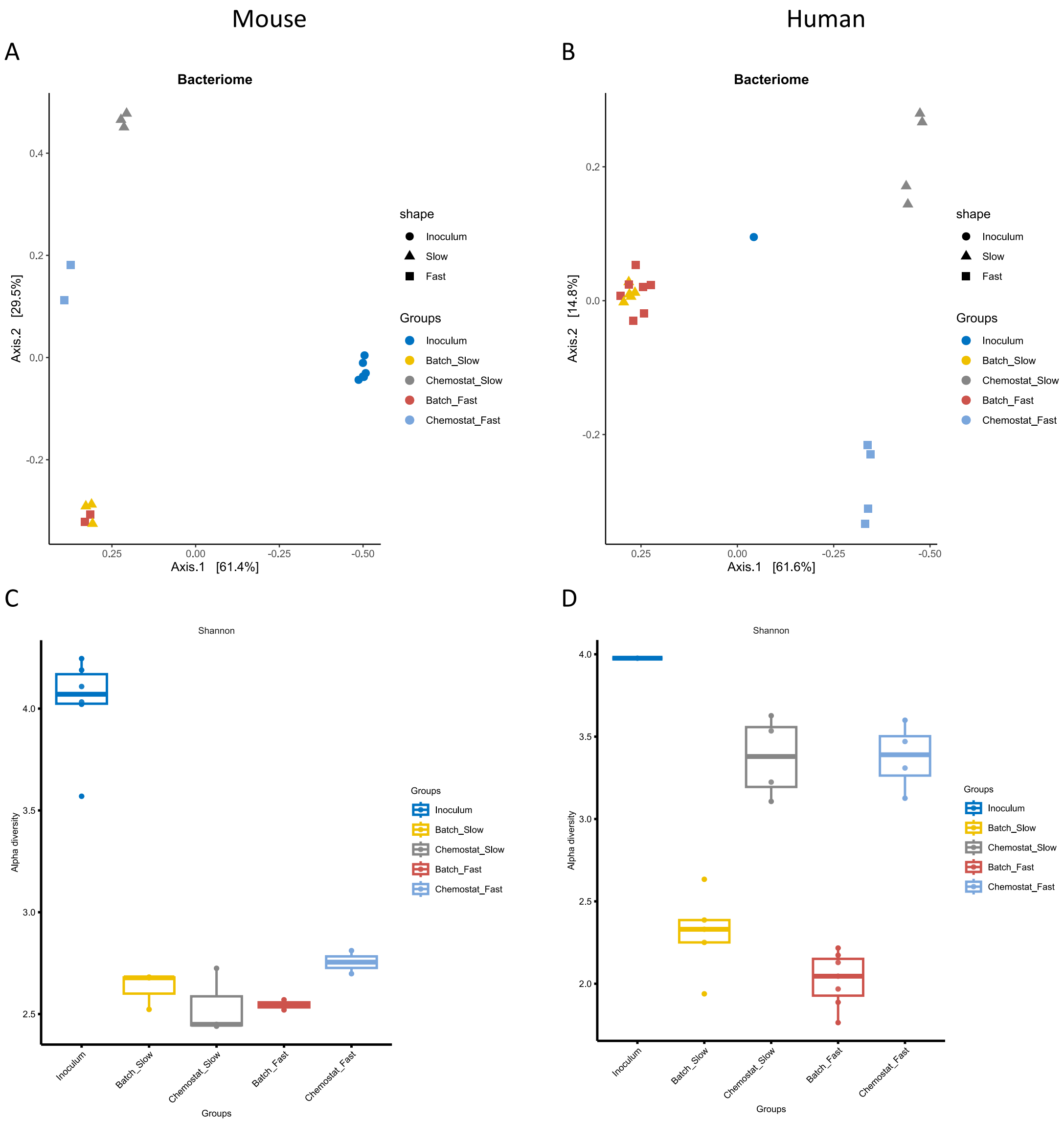


Supplementary Fig. 4. Beta diversity (A, B) and Shannon index (C, D) of bacteriomes of mouse cecal (A, C) and human fecal (B, D) cultures from batch and chemostat cultures. ‘Slow’ and ‘fast’ in the column names indicate the dilution rate used in the chemostat ($D_{low}$ 0.05 and $D_{high}$ 0.2 1/h, respectively).

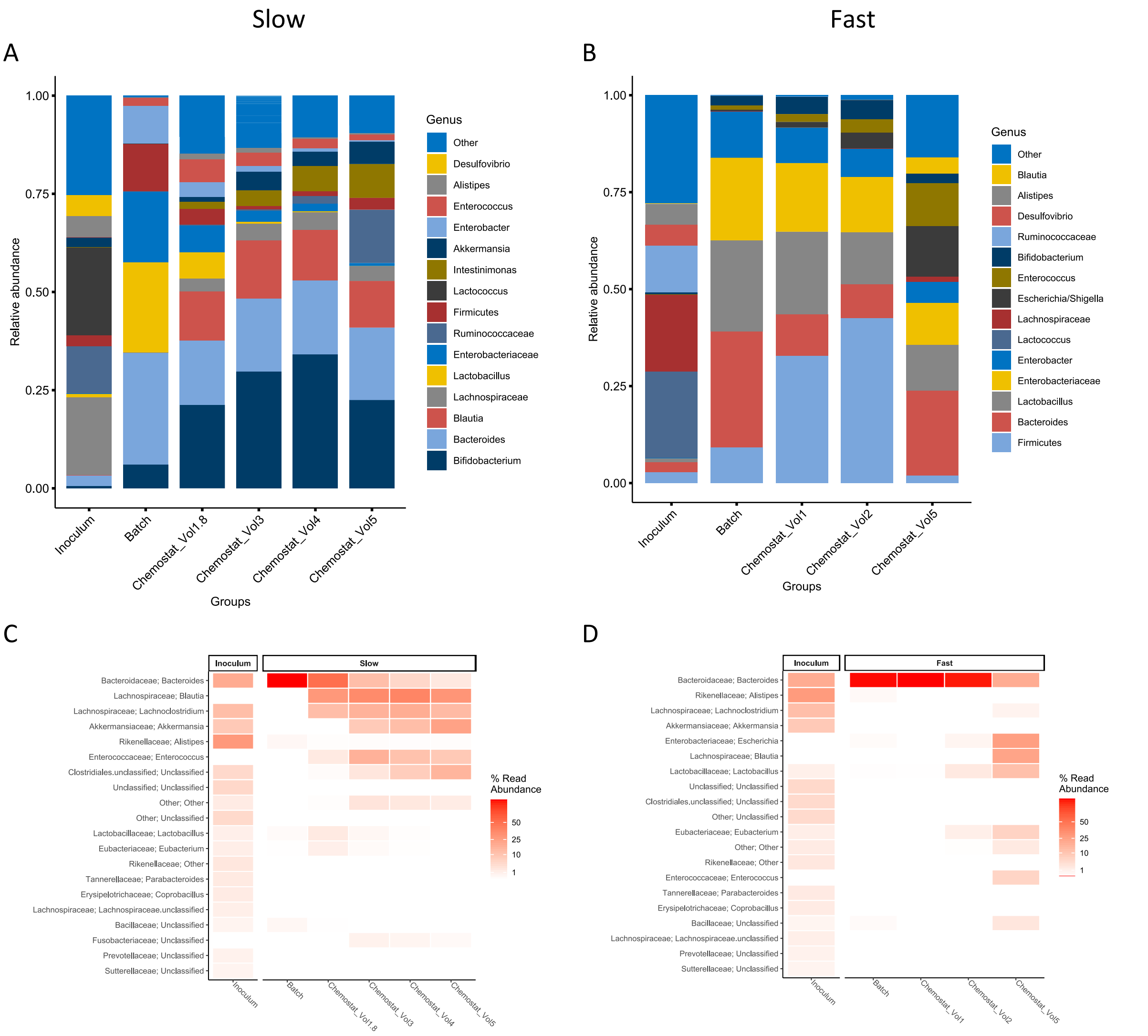


Supplementary Fig. S5. Stabilization of bacteriomes (A, C) and predicted hosts from viromes (B, D) of mouse cecal cultures after switch from batch to chemostat mode. ‘Slow’ and ‘fast’ in the column names indicate the dilution rate used in the chemostat ($D_{low}$ 0.05 and $D_{high}$ 0.2 1/h, respectively). ‘Vol’ indicates the residence times.

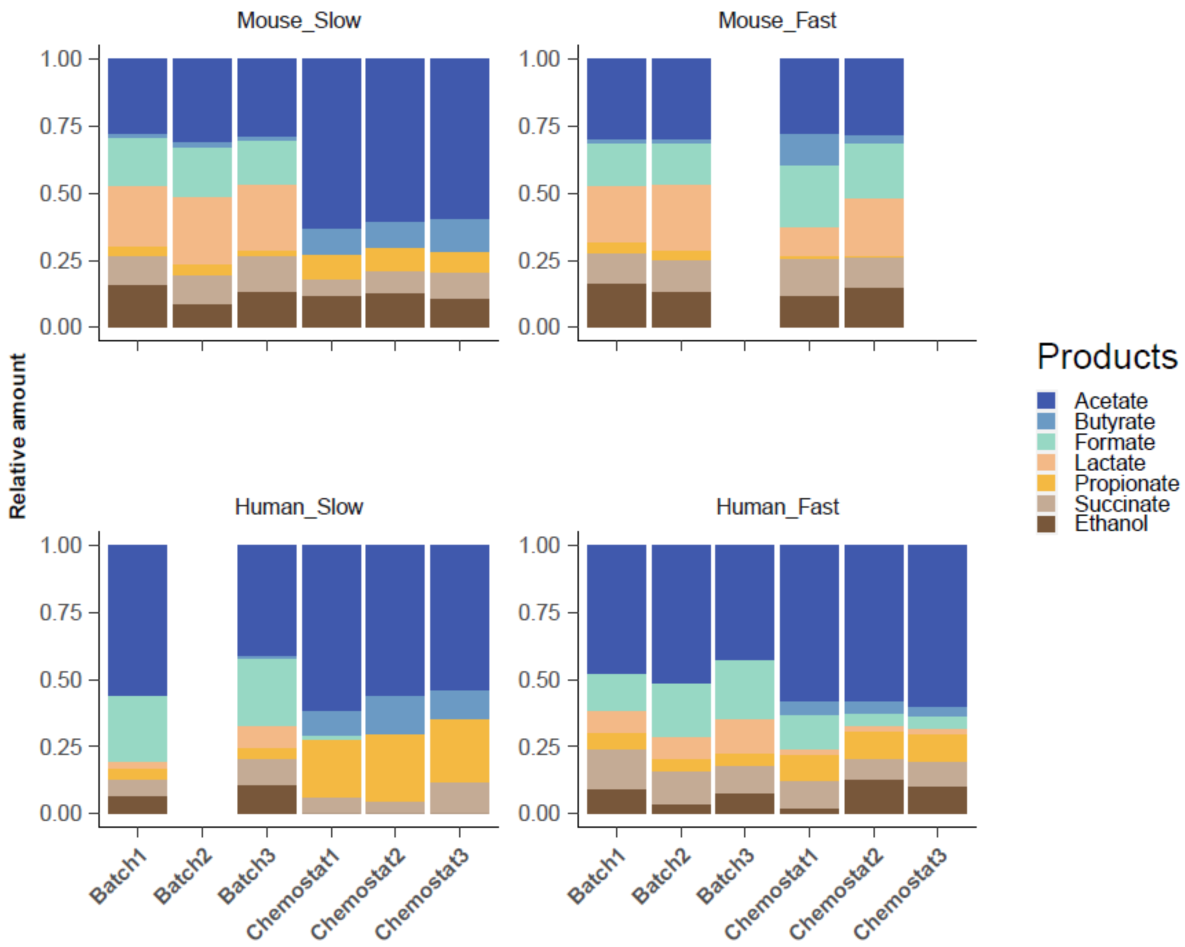


Supplementary Fig. S6. Reproducibility of batch and chemostat cultivations of mouse cecal and human fecal cultures based on metabolite production. Relative amount is presented as the part of a certain product from the sum of all products. The number in the label after ‘Batch’ or ‘Chemostat’ indicates the number of replicate. ‘Slow’ and ‘fast’ in the column names indicate the dilution rate used in the chemostat ($D_{low}$ 0.05 and $D_{high}$ 0.2 1/h, respectively).
